# The Fractional Coalescent

**DOI:** 10.1101/195677

**Authors:** Somayeh Mashayekhi, Peter Beerli

**Affiliations:** Department of Scientific Computing, Florida State University, Tallahassee, FL 32306

## Abstract

A new approach to the coalescent, the *fractional* coalescent (*f*-coalescent), is introduced. Two derivations are presented: first, the *f*-coalescent is based on an extension of the discrete-time Wright-Fisher model. In this extension, for the population of size *N*, the probability that two randomly selected individuals have the same parent in the previous generation depends on the variable α. Second, the *f*-coalescent is based on an extension of the discrete-time Canning population model that the variance of the number of offspring is assumed as a random variable which depends on the variable α. In the second derivation, the *f*-coalescent emerges also as a continuous-time semi-Markov process. The additional parameter *α* affects the variability in the patterns of the waiting times; values of *α* < 1 lead to an increase of short time intervals, but allows occasionally for very long time intervals. When *α* = 1, the *f*-coalescent and Kingman’s *n*-coalescent are equivalent. The mode of the distribution of the time of the most recent common ancestor in the *f*-coalescent is lower than *n*-coalescent when the number of sample size increases, and the time which modes happen on that is smaller compare to *n*-coalescent. Also, this distribution showed that the *f*-coalescent leads to distributions with heavier tails than the *n*-coalescent. Also, the probability that *n* genes descend from *m* ancestral genes for *f*-coalescent is derived. The *f*-coalescent has been implemented in the population genetic model inference software MIGRATE. Simulation studies suggest that it is possible to infer the correct *α* values from data that was generated with known *α* values. When data is simulated using models with *α* < 1 or for three example datasets (H1N1 influenza, Malaria parasites, Humpback whales), Bayes factor comparisons show an improved model fit of the *f*-coalescent over the *n*-coalescent.

---

In 1982, Kingman [1, 2] introduced the *n*-coalescent. The *n*-coalescent describes the probability density function of a genealogy of samples embedded in a population with fixed size. Extensions to this probabilistic description of the genealogical process include changing population size [3, 4], immigration [5, 6], population divergence [7], selection [8], and recombination [9]. These theoretical advances resulted in several widely use computer packages that estimate various population parameters (for example, [10, 11, 12]). While the waiting times for events in the *n*-coalescent are exponentially distributed, a more general framework of these waiting times is offered by the field of fractional calculus. Fractional calculus has attracted considerable interest because of the ability to model complex phenomena such as continuum and statistical mechanics [13], viscoelastic materials [14, 15], and solid mechanics [16]. We introduce Fractional Calculus into population genetics. Our work concentrates on the use of the fractional Poisson process [17] in the context of the coalescent, and we (see the Appendix The Mittag-Leffler Functions) introduce a new model of coalescent, the fractional coalescent or *f*-coalescent. We derive the *f*-coalescent in two independent ways, as an extension of discrete-time Wright-Fisher model and as an extension of the discrete Canning model, and present the properties of the *f*-coalescent. This *f*-coalescent is then implemented in a Bayesian estimator of effective population size; we discuss the implementation and runtime characteristics. We explore the quality of the inference for simulated datasets and also apply the new method to three real data sets: mitochondrial sequence data of humpback whales [18], mitochondrial data of the malaria parasite *Plasmodium falciparum* [19], and complete genome data of the H1N1 influenza virus strain collected in Mexico City in 2014.

## 1 Fractional Coalescent

In this section the fractional coalescent is derived by the extension of discrete-time Wright-Fisher model, and the extension of discrete-time Canning model, and in the Canning model, also the *f*-coalescent emerges as a semi-Markov process in a equivalent way as the *n*-coalescent emerges as a continuous-time Markov process.

### 1.1 The fractional coalescent as an expansion of the discrete-time Wright-Fisher model

In the Wright-Fisher model, the number of offspring of individuals in one generation is given by multinomial distribution [2],

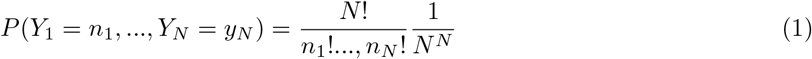

where *N* is the population size and *Y*_*i*_, *i* = 1,…, *N* shows the number of offspring for each individual. When the population size is large, the number of offspring of each individual in one generation approximated by Poisson distribution with mean 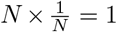. We can interpret this result by using the general form of Poisson distribution [17]

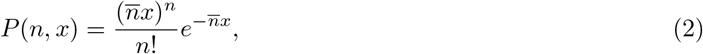

where *P*(*n*, *x*) is the probability of having *n* offspring per generation where we use *x* as the unit time of generation, and the parameter 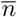 is the average number of offspring occurring per generation. Based on this model the expected number of offspring for each individual is 1.

Looking backward in time, in a haploid Wright-Fisher population of size *N*, the probability that two randomly selected individuals have the same parent in the previous generation is 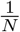, and the probability that they have different parents is 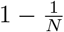. The probability that they do share a common parent after *t* generations is 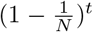. By using a suitable timescale τ such that one unit of scaled time corresponds to *N* generations, the probability that the two lineages remain distinct for more than one unit of scaled time is

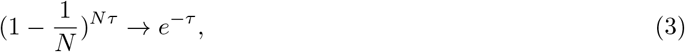

as *N* goes to infinity. Thus, in the limit, the coalescence time for a pair of lineages is exponentially distributed with mean 1 [20]. The *n*-coalescent generalized the two lineages framework to *k* lineages by changing 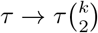, so the probability that the two lineages among *k* lineages remain distinct for more than one unit of scaled time is

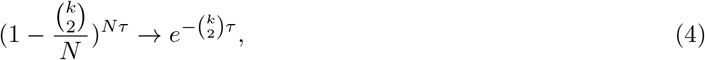

as *N* goes to infinity. When we express τ = *t*/*N* then we recognize the familiar *n*-coalescent formula.

This argument shows, in the *n*-coalescent, the expected number of offspring for each individual is not dependent on the population. Suppose the expected number of offspring for each individual is unknown and is depend on the population. We choose *α* as a unknown value which changes in the different populations, and introduce the unit time of generation and the expected number of offspring based on *α*. Let the number of offspring of each individual in one generation approximated by fractional Poisson distribution where this distribution is similar to Eq. (2) but instead of exponential function we have a summation as

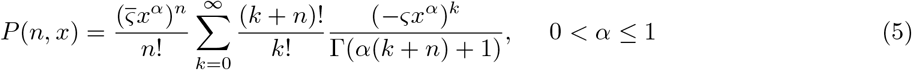

where ς becomes 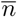 at *α* = 1. In this case *x*^*α*^ is the unit time of generation and the expected number of offspring for each individual is 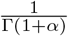 per time of generation when we suppose ς =1.

Looking backward in time, in population, which the expected number of offspring is dependent on *α*, the probability that two randomly selected individuals have the same parent in the previous generation is 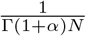, and the probability that they have different parents is 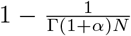. The probability that they do share a common parent after *t*^*α*^ generations is 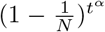. By using a suitable timescale τ^*α*^ such that one unit of scaled time corresponds to *N* generations, the probability that the two lineages remain distinct for more than one unit of scaled time is

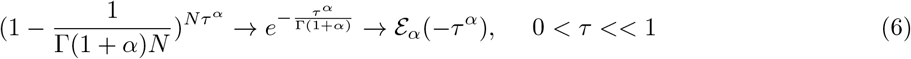

as *N* goes to infinity where 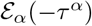 is the Mittag-Leffler function (see Appendix The Mittag-Leffler Functions). Thus, in the limit, the coalescence time for a pair of lineages is distributed as a Mittage-Leffler distribution.

The *n*-coalescent generalized the two lineages framework to *k* lineages by changing 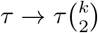. Similarly we can generalized the *f*-coalescent from two lineages framework to *k* lineages by changing 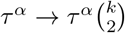.

### 1.2 The fractional coalescent as an expansion of the discrete-time Canning model

The Wright-Fisher model can be generalized. For example, it can be to extended to the Canning model in which the number of offspring of all individuals (*Y*_*i*_) has the same variance (*σ*^2^). This variance changes the time scale to 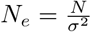 in which

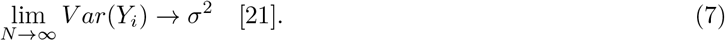

In this case, the probability that the two lineages remain distinct is

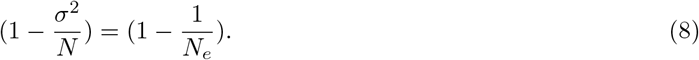

The probability that the two lineages remain distinct for more than one unit of scaled time, which is proportional to *N*_*e*_, is

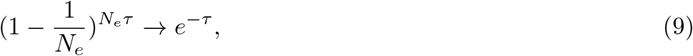

as *N*_*e*_ goes to infinity. [21] has introduced a specific example, the nest-site model, which leads to the Canning model. In this model, the habitat structure determines the distribution of offspring numbers: nests of type *i* comprise a fraction *β*_*i*_ of the total number of nest sites. The quality of nest sites is fixed so that the individuals who occupy sites of type *i* account for a fraction *χ_i_* of offspring. Since in this model *β*_*i*_ and *χ*_*i*_ are fixed, Kingman’s coalescent, by choosing the population size as 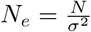 is an appropriate model to capture the common ancestor, where 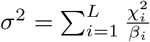 and *L* shows the number of different kinds of nest sites.

If in this model the quality of nest sites is a random variable then Kingman’s coalescent can not be an appropriate model to capture the common ancestor. Suppose *χ*_*i*_ is a random variable whose possible values are 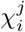, *j* = 1, 2,…. Similar to [21], as *N* increases, the probability of coalescence becomes

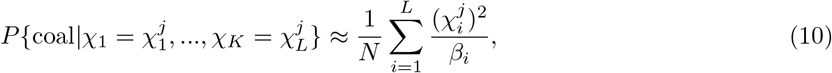

and the variance of the number of offspring *σ*^2^ becomes a random variable whose possible values are 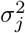, *j* = 1, 2, … where

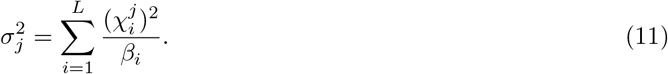

Using Eq. (10) and (11) we have

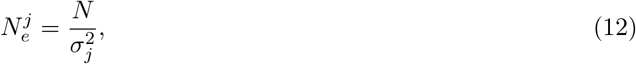

and

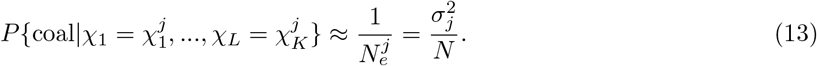

Assume that the variance of number of offspring is distributed as a random variable (*σ*^2^ ∈ (0, ∞)) that has the probability density function ω(*σ*^2^, *α*) where 0 < *α* ≤ 1; this random variable depends on the population. By this assumption, similar to Eq. (8) the probability that the two lineages remain distinct for more than one unit of scaled time is

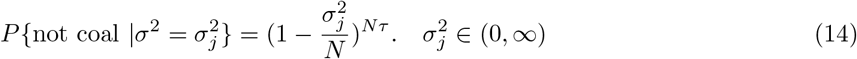

The average of Eq. (14) over the distribution of *σ*^2^ ∈ (0, ∞) shows the probability that the two lineages remain distinct for more than one unit of scaled time as

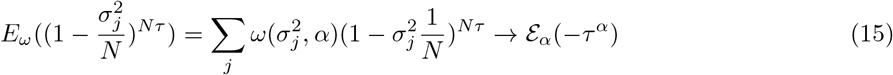

as *N* goes to infinity, 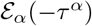 is the Mittag-Leffler function (see Appendix The Mittag-Leffler Functions). We choose the time scale as 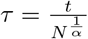;

Thus, in the limit, the coalescence time for a pair of lineages is distributed as the fractional generalization of the exponential distribution [22]. Similar to the discussion which we had in the last section we can generalized the *f*-coalescent from two the lineages framework to *k* lineages by changing 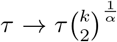, and in this case the probability that the two lineages among *k* lineages remain distinct for more than one unit of scaled time is

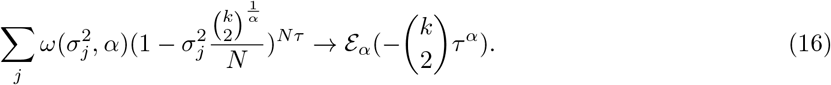

Choosing the time scale as 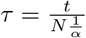 keeps the parameter the same as the *n*-coalescent.

### 1.3 *f*-coalescent as a semi-Markov process

We can, also, describe the *n*-coalescent as a Markov process. We introduce a stochastic matrix *P*_*N*_ which is a single-step transition matrix as

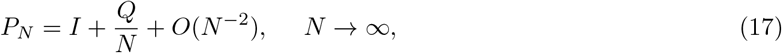

where *Q* is the rate matrix of a Markov-process. With an appropriate time scale proportional to *N*, after *t* generation we have

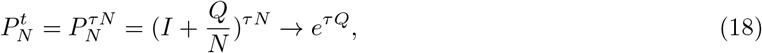

as *N* goes to infinity. Eq. (18) is similar to Eq. (4) but in matrix form.

Using Eq. (16), we can find the following form of a Mittag-Leffler matrix function

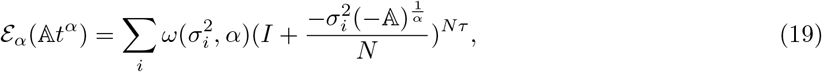

where 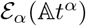 is a stochastic matrix, and matrix 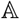 has two properties: Off-diagonal entries are positive or zero (*a*_*ij*_ ≥ 0, *i* ≠ *j*) and diagonal entries ensure rows sum are zero (*a*_*ii*_ = − ∑_*j*, *j*≠*i*_ *a*_*ij*_). In the discrete case, the variance of the number of offspring can be a random variable, *σ*^2^. For all 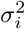 we need to introduce a single-step transition matrix. The rate matrix *Q* has the same properties as matrix 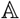. In Eq. (17), we substitute *Q* with 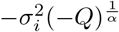 and rewrite *P*_*N*_ as

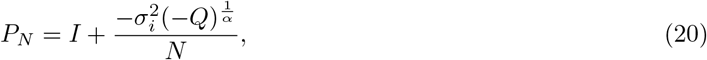

and by introducing the appropriate time scale which is proportional to *N*, after *t* generation we have

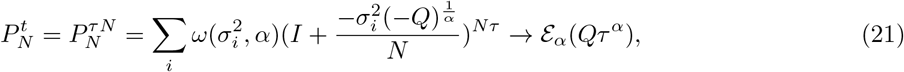

Eq. (21) is similar to Eq. (16) but in matrix form. We can interpret 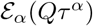 as a weighted sum of related Markov processes which itself is a semi-Markov process (see Appendix The Mittag-Leffler Functions).

These derivations of the *f*-coalescent from the Wright-Fisher model and Canning model suggest that we have a versatile framework that maintains the strictly bifurcating property of the *n*-coalescent, but permits more variability in waiting times between coalescent events. Thus, this versatility may allow to infer processes that change the waiting times, for example selection, better. Currently, multi-furcating coalescent, such as the Bolthausen-Sznitman-coalescent [cf. 23], are used to discuss such forces. The *f*-coalescent may be a viable alternative.

### 1.4 Time to the most recent common ancestor (TMRCA)

The *n*-coalescent has two steps: first, choosing a pair to coalesce using the concept of equivalence classes. Second, the waiting time in which two lineages need to coalesce. For the *f*-coalescent, we changed the second step resulting in a different TMRCA than for the *n*-coalescent. We derive this new distribution of the TMRCA of the *f*-coalescent, and compare it with the TMRCA of the *n*-coalescent. We also present the probability that *n* genes are descendants from *m* ancestral genes using the *f*-coalescent, and compare these results with the *n*-coalescent. To do this, we extend the work of [24] to the *f*-coalescent. In the following Theorems we use Eq. (6), (15) and (A6). These lead to the probability density of waiting time in the *f*-coalescent

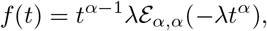

where for *α* = 1 we have the exponential distribution which has been used in the *n*-coalescent.

#### Theorem 1

**If** 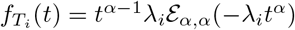 **is the distribution of a waiting time in the *f*-coalescent where** *T*_*i*_, *i* = 2, …, *n* **are the coalescent times and** 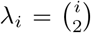 **then the distribution of** 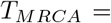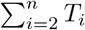 **is as follows**

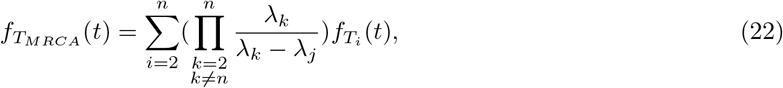

**or, equivalently, this can be presented as**

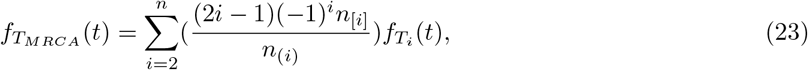

**where**

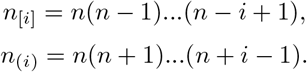

**Proof: Please see the Appendix**.

**Conclusion**: We have plotted Eq. (22) (or Eq. (23)) in Figure 1 for different values of *α*. *α* = 1 is related to the *n*-coalescent. For small value of *α* (*α* = 0.3) the distributions are the same for all value of *n*. This result will be confirmed in the section Methods where we see the datasets generated with *α* < 0.4 most commonly did not have any variable sites. For 0.3 < *α* < 1, the mode of 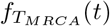 is decreased when *n* is increased, and modes are about the same point for all value of *n* (The mode of 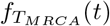 moves very slowly to the right). For *n*-coalescent (*α* = 1) the mode of 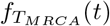 is fixed but they are at different points for different values of *n* (The mode of 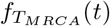 moves fast to the right, but the value of mode does not change as *n* increase). The asymmetry of 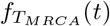 is stronger for *α* < 1. While for *α* = 1 as *n* increase, the distribution of 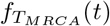 convergence on a distribution with mean equal 2 which corresponds to a period 2*N* generations under the haploid Wright-Fisher model [21], for *α* < 1 we have a heavy-tailed distributions as *n* increases.

**Figure 1:**
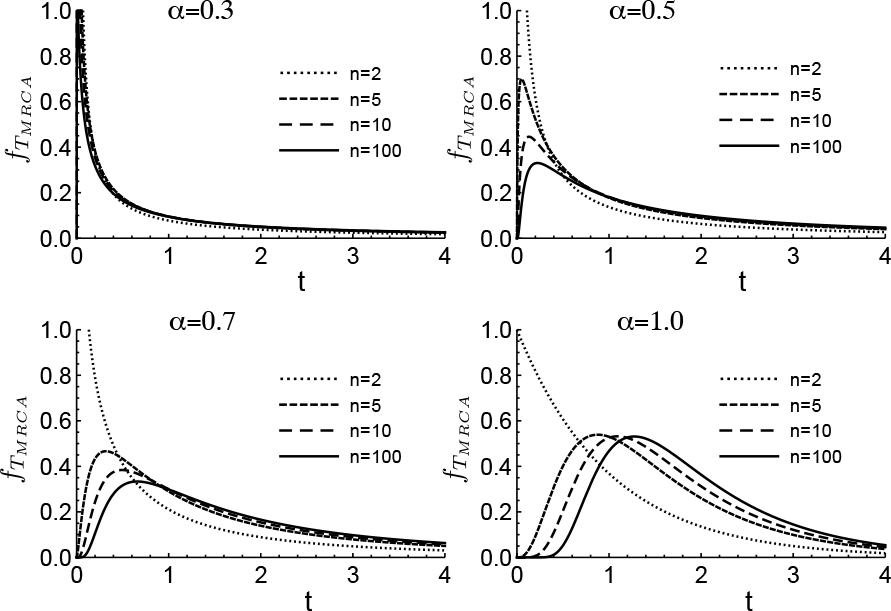
Distribution of the time to the most recent common ancestor for different value of *α*

In the next theorem, we derive the probability that *n* genes are descendants from *m* ancestral genes in the *f*-coalescent.

#### Theorem 2

**In the *f*-coalescent, the probability** *P*_*nm*_(*T*) **that** *n* **genes are descended from** *m* **genes** *T* **units of time ago is**

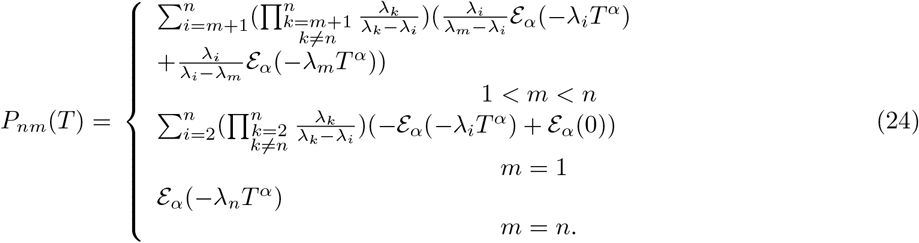

**Proof: Please see the Appendix**.

Conclusion: Since for *α* < 1, *E*_*α*_(−λ_*i*_*T*^α^) is greater than 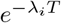, in the *f*-coalescent as *T* increases *m* does not decrease as rapidly as in the *n*-coalescent [24]. We give some numerical value of Eq. (24) for different value of *α* in the Supplementary Informations (Numerical values of the probability that *n* genes are descended from *m* ancestral). In these tables *α* = 1 is the *n*-coalescent which in this case *m* decreases quite rapidly as *T* increase, but this is not the case in *f*-coalescent.

It should be noted that we derive the time to the most recent common ancestor in section Methods empirically too, to compare *f*-coalescent with BS-coalescent.

### 1.5 Probability density function of a genealogy based on the *f*-coalescent

The waiting times in Kingman’s *n*-coalescent have an exponential distribution and we express the probability density function of a time interval *u*_*k*_ with *k* lineages as

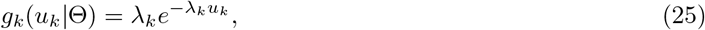

where

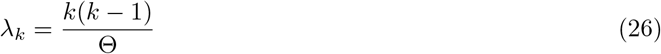

with the mutation-scaled population size Θ = 2*N μ* where *N* is effective population size and *μ* is mutation rate per generation. For the *f*-coalescent we express the waiting time with the fractional generalization of the exponential probability distribution. The exponential function is replaced by the generalized Mittag-Leffler function 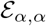 which has an additional parameter *α* (The Mittag-Leffler Functions). In this section we follow the formulation which we derived in the extension of Canning model, but we can have a similar discussion for the Wright-Fisher model. Using Eq. (16), the probability that two of *k* lineages coalesce after one unit of scaled time is

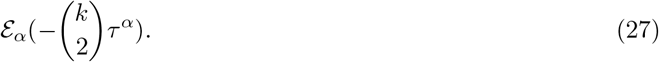

When we replace the scaled time τ with 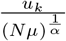, where *u*_*k*_ has been scaled based on mutation 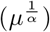, we can rewrite Eq. (27) as

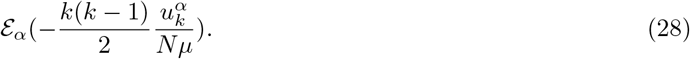

Replacing 2*N μ* with Θ we get

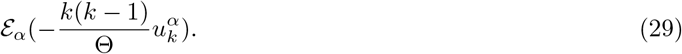

Using Eq. (29) and (A6), the probability density for a time interval *u*_*k*_ is

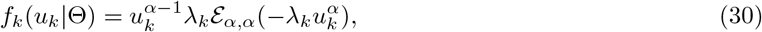

where λ_*k*_ is introduce in Eq. (26). We consider values for *α* in the interval (0,1]. The Mittag-Leffler function reduces to the exponential function with *α* =1 and Eq. (30) reduces to the familiar probability density for a time interval *u*_*k*_ for *n*-coalescent. The probability density of all observed *u*_*k*_ with *k* = 2, …,*K* given the mutation-scaled population size Θ is the joint probability distribution function as

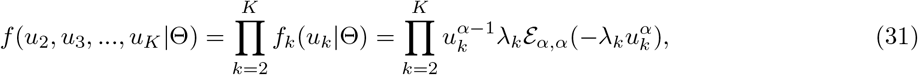

where *K* is the number of samples. To extract a particular genealogy *G* out of the many possible topologies defined by the interval times *u*_2_, *u*_3_,…, *u*_*K*_ we need to take into account the number of possible configurations at any time *u*_*k*_; using Kingman’s *n*-coalescent for any *u*_*k*_ and *k* lineages there are 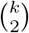 potential configurations and we pick one particular one. If we use the same assumption for *f*-coalescent that only two lineages per generation can coalesce then we get

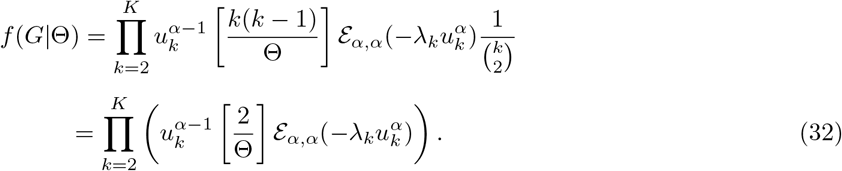

### 1.6 Implementation

We implemented our method in the existing framework of the software MIGRATE [10]. In this framework we approximate the Bayesian posterior density *f*(*ρ*|*X*, *α*) = *f*(*ρ*)*f*(*X*|*ρ*, *α*)/*f*(*X*|*α*) where *X* is the data, ρ is the parameter set for a particular population model, and *α* is a fixed parameter for the Mittag-Leffler function. The software uses MCMC to approximate the posterior density, calculating *f*(*ρ*) and *f*(*X*|*ρ*, *α*). To choose tree genealogy during the MCMC we draw new times for events. Details of the tree changing algorithm are described by [25]. To draw a new time (*t*_0_), we solve

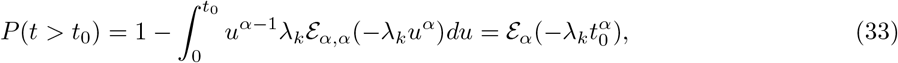

where *k* and *α* are fixed numbers and we choose a random numbers between (0,1) for *P*(*t* > *t*_0_). Since using Eq. (33) directly to draw times is time-consuming, we use the sampling method of the Mittag-Leffler distribution which has been introduced by MacNamara [22].

Since the Mittag-Leffler function can be expressed as a mixture of exponential functions, the fast simulation of geometric stable distributions can be used to sample the time. In this method, the time drawing from Mittag-Leffler distribution is multiplication of random numbers which are distributed by *ω*(*κ*, 1) and the exponential distribution. As a result, the time derived from the Mittag-Leffler function is

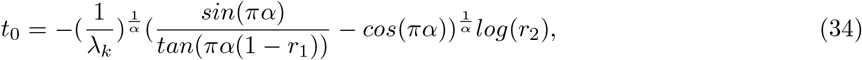

where *r*_1_ and *r*_2_ are two independent random numbers. More details for Eq. (34) has been presented by MacNamara [22].

A particular problem for a useful implementation was the calculation time of the generalized Mittag-Leffler function. We programmed the function in the C-language using the algorithm described by Gorenflo [26] that also was used to create a Matlab-function by I. Podlubny (https://www.mathworks.com/matlabcentral/fileexchange/8738-mittag-leffler-function). We compared our C Mittag-Leffler function with the one implemented in Mathematica 11.0.1.0 [27] for correctness. During these tests we realized that our implementation will be very slow in a MCMC run, because the function will be called millions of times. We implemented a lookup table for values of The Mittag-Leffler Functions 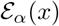 and 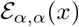 for *α* ∈ [0.01,0.02, 0.03, …0.99]. The lookup-table values were created by finding all values for a given α that have 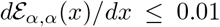 and 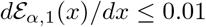. This lookup-table is then used with a Hermite-cubic-interpolation and a linear interpolation at the table end points to return values for The Mittag-Leffler Functions. This interpolation is roughly 2x faster than our original C-function. Our lookup table and interpolation function can be downloaded from http://popgen.sc.fsu.edu/codes.

## 2 Methods

*Time to the most recent common ancestor for the f-coalescent*:The most recent common ancestor of the sampled individuals is at root of a genealogy. The time of this most recent common ancestor (TMRCA) is directly related to the effective population size of the population which the sample was taken from it. Coalescence theory assume that each time interval *j* is independent from the time interval *j* − 1 or *j* + 1. If we are only interested in the population size, we only need to consider the time intervals and do not need to know details of the topology of the genealogy. We compared empirical distributions of the TMRCA for a sample of 5 individuals for the *f*-coalescent, *n*-coalescent, and Bolthausen-Sznitman-coalescent [BS-coalescent; 28]. BS-coalescent allows merging of more than two lineages and is used to describe the waiting times in a genealogy that is affected by selection [23]. All calculations were executed in Mathematica 11.0.1 [27]. For the *n*-coalescent, BS-coalescent, and the *f*-coalescent we summed independently drawn time intervals for 5, 4, 3, and 2 lineages to get the TMRCA. We drew 100,000 replicated TMRCAs for each of the three *n*-coalescent settings (strong population shrinkage, fixed population size, strong population growth), two BS-coalescent settings (*T*_*c*_ = 0.005 and 0.01) and the two *f*-coalescent settings (*α* = 0.9 and 0.8). For the fixed population size *n*-coalescent time intervals, we solved Eq. (3) in [25]; for time intervals of the growing and shrinking population we used Eq. (4.6) in [29]; for the BS-coalescent we used the algorithm of 23]; for the *f*-coalescent we used Eq. (34).

*Simulated data*:We evaluated the algorithms using simulations. We updated our simulator package (available at peterbeerli.com/software and bitbucket.com) to allow generating genealogies from the *f*-coalescent. We generated 100-locus datasets with a population size *N*_*e*_ = 2500 and mutation rate per site per generation *μ* = 0.000001 for *α* in [0.4,0.5, ‥1.0]. Each locus had 10,000 base pairs. Datasets generated with *α* < 0.4 most commonly did not have any variable sites. The datasets were then analyzed with MIGRATE with a fixed *α* of 0.4 or 0.5,…, or 1.0. Each dataset was thus analyzed 7 times with different *α* values allowing us to compare the different runs considering each *α* value as a different model. We use the framework of marginal likelihoods [cf. 30] to compare the different models. The method implemented in MIGRATE uses thermodynamic integration to approximate the marginal likelihoods [31, 32]. We then used these marginal likelihood as model weights [33] to decide whether models could be considered to be appropriate for the data or not.

The *f*-coalescent has an additional parameter *α* which could reflect hidden structure. To explore whether the parameter *α* responds to population structure, we simulated data using the structured coalescent for two populations exchanging migrants using three different magnitudes of gene flow: assuming only 1 migrant per 100 generations, 1 migrant per 4 generations, and 10 migrants per generation. These data were then analyzed with the structured *n*-coalescent, the non-structured *n*-coalescent and non-structured *f*-coalescent. We then compared the marginal likelihoods of these different models. We reanalyzed this same data focusing only on population 1 assuming that we did not recognize population 2, using single population models for the *n*-coalescent and the *f*-coalescent, and a ghost-population model [34] that assumes we know about a second population but do not have any data for it.

*Real data*:We used three biological datasets to explore whether the *f*-coalescent could be a better fit than Kingman’s *n*-coalescent. The first dataset is a dataset of short mitochondrial control region sequences of the humpback whale population in the North Atlantic [18] (H data), the second is a small dataset of the malaria parasite *Plasmodium falciparum* in Africa [19] (P data), and the third dataset is a small dataset of the H1N1 influenza strain the scared the world in 2014 with an outbreak in Mexico (I data) (downloaded on August 7 2017 from www.fludb.org). We used a simple mutation model, F84 + G [35, 36], optimized the transition/transversion ratio (ttr) and the site rate parameters in paup* [37]. For the whale and Plasmodium data the ttr were 18.08 and 1.56, respectively. The site rate variation parameter estimates were 0.0277 and 0.00982. The influenza data were heterogeneous: segment 1-4 had a site rate variation parameter of ~ 0.02, and the segments 5-8 had a value of > 200.0. We analyzed the total influenza data as 8 independent loci using the F84 model. All data were then run using *α* = [0.4,0.5,…, 1.0]. The influenza data was also analyzed using a exponential growth *n*-coalescent model. All models were then compared using marginal likelihoods.

## 3 Results

*Time to most recent ancestor for f-coalescent*: Figure 2 shows examples of empirical distributions of the TMRCA for a sample of five individuals for the *n*-coalescent, the *f*-coalescent with two different values for *α* and BS-coalescent. Each curve is based on 100,000 simulated TMRCA values. With *α* < 1 the distribution becomes more peaked with more shorter time intervals and rare, but large time intervals, that lead to heavier tails than with the *n*-coalescent; Median values for the TMRCAs of the different models were: 0.00379 for *f*-coalescent with *α* = 0.9 and 0.00026 with *α* = 0.8; 0.00667, 0.00343, 0.00031 for *n*-coalescent with no growth, strong growth, and strong shrinkage, respectively; BS-coalescent had medians of 0.00576 with *T*_*C*_ = 0.01 and 0.00288 with *T*_*C*_ = 0.005. The expectation of the TMRCA for the *n*-coalescent for 5 samples simulated with a Θ = 0.01 is 0.008. The expected TMRCA for the *f*-coalescent is infinite because of the heavy tail [cf. 38]. Comparison with the BS-coalescent are more difficult because of the parametrization. Neher [23] postulates that the coalescent rates of the BS-coalescent and the *n*-coalescent are not equivalent, the parameter *T*_*c*_ in BS-coalescent defines the time to the selectively positive innovation whereas in the *f*-coalescent and the *n*-coalescent the rate is defined by the scaled population size. There is certainly a direct relationship between *T*_*c*_ and Θ but under selection that relationship changes. We recognize that distributions of some the *f*-coalescent TMRCA and some of the BS-coalescent TMRCA look rather similar compared to the others, but the mapping of the parameter *T*_*c*_ and Θ will need further investigation.

**Figure 2:**
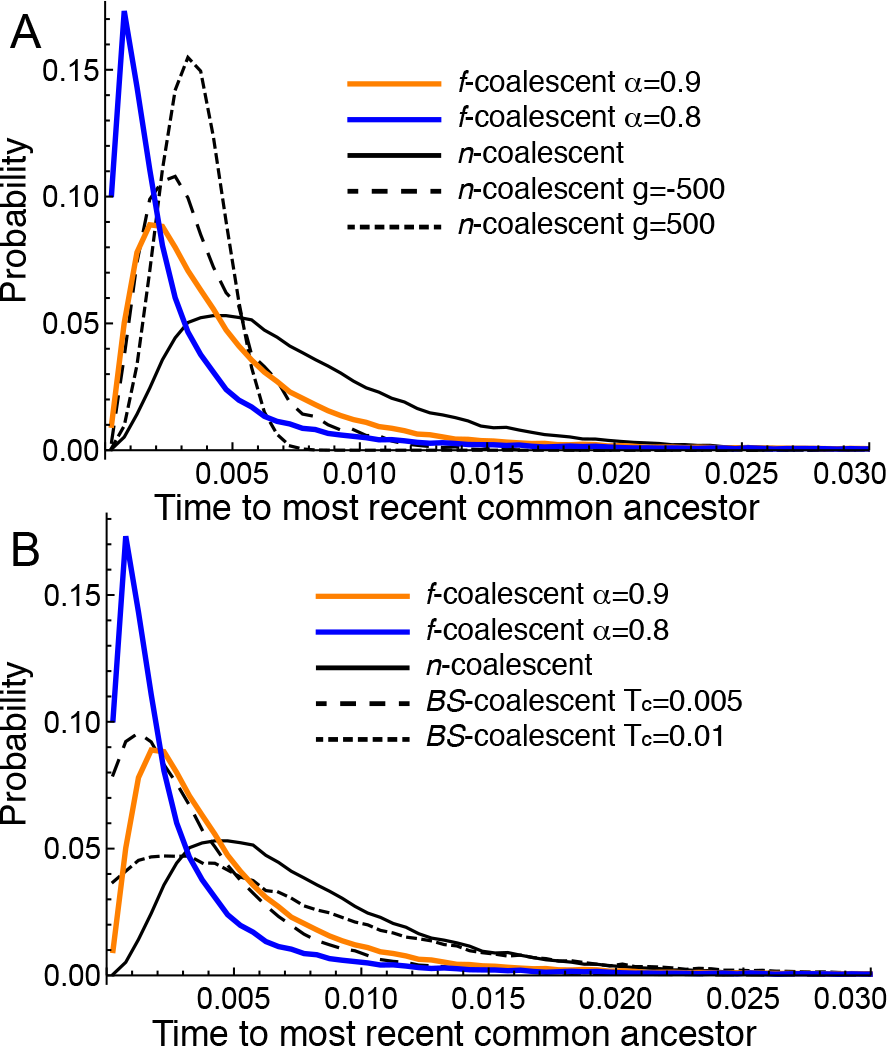
Empirical distribution of the time of the most recent common ancestor for various coalescents: (A) strictly bifurcating; (B) *f*-coalescent versus *n*-coalescent and multifurcating Bolthausen-Sznitman coalescent. The x-axis is truncated at 0.03. Each curve represent a histogram of 100,000 draws of the TMRCA.

*Simulation*: Figure 3 shows that it will be difficult to recover the parameter *α* that was used to simulate the data; data simulated with a particular *α* has a considerable range when estimated. A problem with these simulations is that the number of variable sites is dependent on the mutation-scaled population size Θ and the simulated branch length. Despite of having the same Θ = 0.01 among all simulations, the number of variables sites varies considerably over the range of *α*: with an *α* = 0.4 about 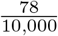 sites (mean) are variable compared to *α* = 1.0 with 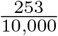 sites. With *α* ≤ 0.4 it is common to see no variable sites in a locus, median among 100 loci is zero. Low variability data sets are difficult to use in any coalescent framework.

*Comparing f-coalescent with structured coalescence and population growth*: Simulated data from a structured coalescence model with two subpopulations exchanging migrants was evaluated under two different scenarios: (A) we sampled data from both populations and (B) we collected data only from population 1. For (A) we ran models that assume that the population is (1) not structured with *α* < 1, (2) the standard single-population *n*-coalescent, and (3) a structured coalescence model. A Bayesian model selection approach excludes all models with *α* < 1. The non-structured standard *n*-coalescent model is rejected for the low and medium gene flow scenarios, but the single-population *n*-coalescent models is the best model for data simulated with high immigration rates; see electronic supplement for details. For (B) we ran models that assume that the population is (1) not structured with *α* < 1, (2) the standard single-population *n*-coalescent and (3) a model that assumes a ghost population [34]. Models that include high to very high immigration picked the *n*-coalescent model as the best, rejecting the *f*-coalescent and the ghost-population model. With low immigration rates the *f*-coalescent model had higher marginal likelihoods at *α* = 0.9 suggesting that rare migrants will disturb the exponential waiting time pattern of a single *n*-coalescent population.

**Figure 3:**
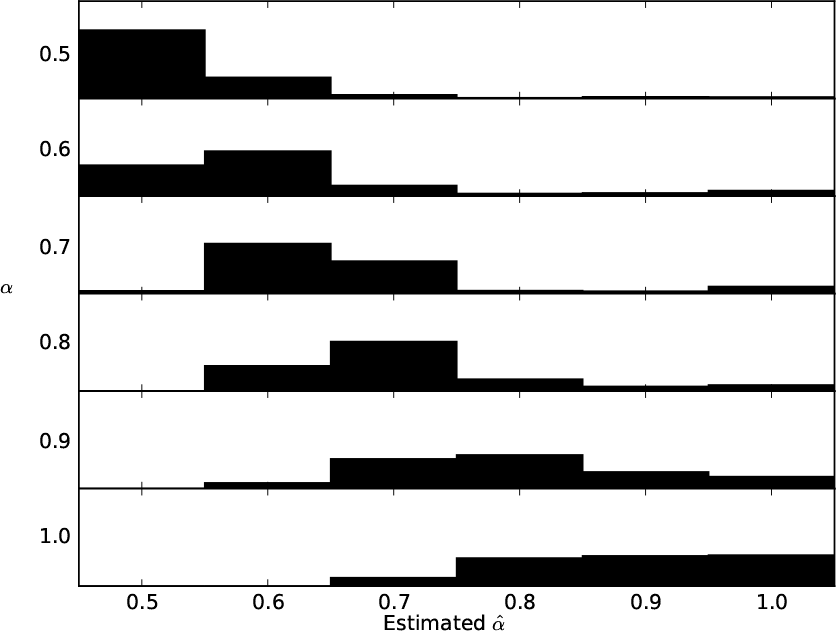
Comparison of recovery of the model under which the data was simulated. *α* was used to simulate the data; The histograms of 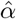 show the number of wins in a marginal likelihood comparison. For each true *α*, 500 simulations with 10 individuals and 10,000 base pairs were used.

*Real data results*: The analysis of the Malaria (P data) and the humpback whale (H data) datasets revealed that the Mittag-Leffler tuning parameter *α* may have a wide range (Fig. 4). The H data had only 283 sites thus parameter reconstruction is not very robust. In contrast, the P data has 5946 base pairs and should allow good reconstruction of the parameters. For P data, model selection using marginal likelihood suggests that models within the range of 0.55 < *α* ≤ 1.0 are preferred; models with an *α* < 0.55 had a model probability of 0.0000. The maximum marginal likelihood was at *α* = 0.8. The estimated mutation-scaled population size, Θ, varied considerably in this range. At *α* = 0.55, Θ was 0.04810 and at *α* = 1.0, Θ was 0.00197. The best model had a mode equal 0.01157 and 95%-credibility for Θ from 0.00687 to 0.01833. Both, the P and the H dataset suggest that the *n*-coalescent model was a good fit. The 8-segment dataset of the H1N1 strain of influenza from Mexico in 2014 had a well-defined maximum marginal likelihood at *α* = 0.85. The influenza dataset was the only dataset among the tree tested dataset that potentially excluded the *n*-coalescent as a reasonable model. Model probability for *n*-coalescent (*α* = 1.0) was 0.0027. We also ran a model that used the *n*-coalescent and exponential growth, estimating two parameters (growth *g* and Θ). The marginal likelihood for the *θ* + *g* model (ln marginal likelihood = ln mL = -19455.27) is lower than the best model with *α* = 0.80 (ln mL= -19338.28) and also lower than the constant-size *n*-coalescent model (ln mL= -19342.24), the ln Bayes factor comparison with the best model was -118.14, suggesting that the growth model is inferior to the *f*-coalescent. Migrate output files are available in the github repository.

## 4 Discussion

A feature of the *f*-coalescent is the ability to accommodate very variable time intervals. Mixtures of very short branch lengths with very large branch lengths are possible, whereas the *n*-coalescent forces a more even distribution of these time intervals. Extensions of the *n*-coalescent to allow for population growth or population structure do not match the variability of time intervals of the *f*-coalescent. With exponential population growth, time interval near the sampling date are enlarged and near the root of the genealogy the time intervals are shortened; the *n*-coalescent with exponentially shrinking populations also have heavy tails, but seem to have more longer branches than the *f*-coalescent. In the *f*-coalescent time interval near the tips are shortened and time intervals near the root are lengthened.

Analyses of data that was simulated using a structured *n*-coalescent model show that only when we remove half of the simulated data and analyze only a single subpopulation with models that assume that this is an isolated panmictic population, we will get a better model fit with a *f*-coalescent model when the immigration rate is 1 per 10 generations. The unique mix of short and long waiting times of the *f*-coalescent thus may allow inferences with unknown compartmentalization; but we will need to extend our single population *f*-coalescent to structured populations to study these type of models.

**Figure 4:**
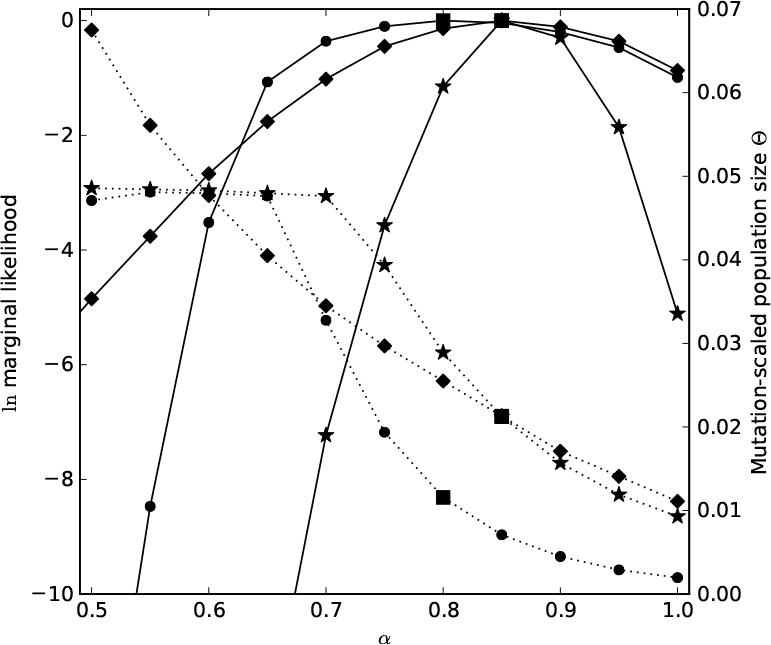
Model selection using relative marginal likelihoods of a *Plasmodium falciparum (circle)*, a H1N1 influenza (star), and a Humpback whale mtDNA (diamond) dataset. The square marks the value of *α* and the estimate for scaled Θ of the best model for each dataset.

The three real data examples suggest that the *f*-coalescent is a better fit to the data (in all of the test cases). Whale dataset consists only of a single population. Roman and Palumbi [18] stated that the immigration rate from the South Atlantic into the North Atlantic population is minimal. The marginal likelihood comparison of different *α* for the Humpback whale, the Plasmodium data did not reject the *n*-coalescent. The influenza dataset rejected the *n*-coalescent barely. This may indicate that the *f*-coalescent may improve our understanding of mitochondrial evolution and fast-evolving organisms that are under selection. We believe that the *f*-coalescent can serve as a model to approach loci under selection, where the positively selected locus will sweep quickly through the population and result in many short coalescent intervals.

The *f*-coalescent is a promising extension of the coalescent because it allows considering a wider range of waiting times between events, but certainly will need more study.

## 5 Acknowledgements

We thank John Wakeley for giving us many suggestions that helped us to improve our manuscript. This work has been supported by the National Science Foundation award DBI 1564822, and the IT resources at the research computing center of Florida State University.

## 6 Appendix

## 6.1 The Fractional Poisson Process

The classical Poisson process, is a common random process in probability theory. Several probability distributions arise from the classical Poisson process, for example the Poisson distribution and the exponential distribution. In this process, the number of events, *N*(*u*), which happen during the time interval from 0 to *u* has a Poisson distribution; and, the waiting times between these events are exponentially distributed. Kingman’s *n*-coalescent uses the classical Poisson process to model the waiting time between two coalescent events. For the *f*-coalescent, we change the distribution of waiting times from an exponential distribution to the fractional generalization of the exponential probability distribution.

Before we introduce the fractional generalization of the exponential probability distribution, we first introduce the fractional Poisson Process.

Fractional Poisson processes have been developed by using the classical Poisson process in two different cases, the time-fractional Poisson process and space-fractional Poisson process [39]. The combination of these two different cases has been introduced as the space-time-fractional Poisson process [39].

For the classical homogeneous Poisson process we know that

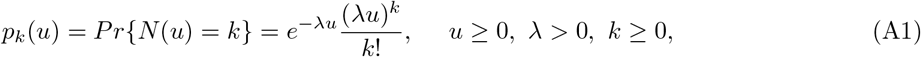

it is also known that *p*_*k*_ (*u*) with *k* ≥ 0 solves the following Cauchy problem

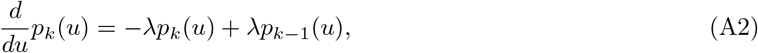

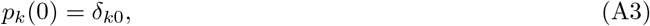

where δ_*n*, 0_ is the Kronecker symbol. Time-fractional generalizations of the Poisson process are based on the substitution of the integer-order derivative operator appearing in Eq. (A2) with a fractional-order derivative operator such as the Riemann-Liouville fractional derivative [17] or the Caputo fractional derivative [40] (see Definitions of Fractional Derivatives). Properties of the time-fractional Poisson process have been studied in [41] and [42]. Space-fractional generalizations of the Poisson process are based on the substitution of the backward shift operator space in Eq. (A2) with the fractional backward shift operator [39].

We use the time-fractional generalization of the homogeneous Poisson process which has been introduced in [17]. In this generalization, the fractional Poisson process has been introduced as the counting process with probability *P*_*α*_(*n*, *u*) of having *n* items (*n* = 0,1, 2,…) during time interval *u* by the following special form of the fractional Kolmogorov-Feller equation for 0 < *α* ≤ 1

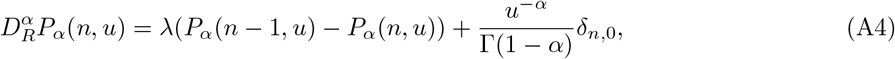

where δ_*n*, 0_ is the Kronecker symbol with normalization condition

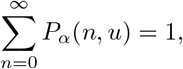

By solving Eq. (A4) we have [17]

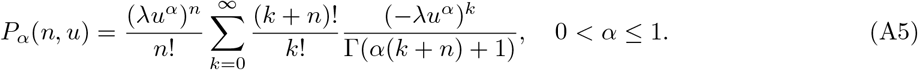

In this generalization, the probability distribution function of waiting time for the fractional Poisson process can be expressed as

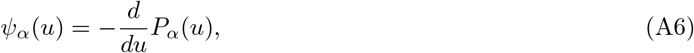

where *P*_*α*_(*u*) is the probability that a given inter arrival time is greater or equal to *u*

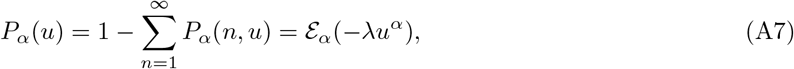

where 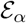 is the Mittag-Leffler function see (The Mittag-Leffler Functions). By using Eqs. (A6) and (A7) the waiting time for fractional Poisson process has the following probability distribution function

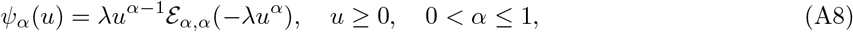

where 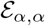 is the generalization of the Mittag-Leffler function (see The Mittag-Leffler Functions). The *ψ*_*α*_(*u*) defined by Eq. (A8) is fractional generalization of the well known exponential probability distribution.

**Figure 5:**
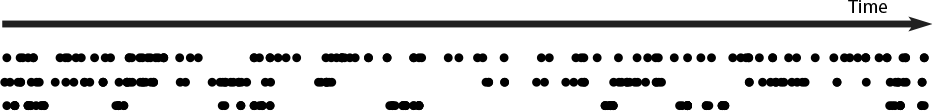
Times between events drawn using the classical Poisson process and the fractional Poisson process with *α* = 0.9, and 0.8; from top to bottom.

The generalization of the Mittag-Leffler function has two different parameters *α* and *β* (see The Mittag-Leffler Functions), and as it has been mentioned in Eq. (A8) the fractional Poisson process, has been derived by using the Mittag-Leffler function using (*α* = *β*). Figure 5 shows the time of 100 independent events which happen in three cases of the Poisson process with various *α* (1.0, 0.9, and 0.8). For calculating these times we have supposed that the probability each event occurs is a random number while λ has been chosen to be fixed and same in all three different cases. With an *α* < 1 more times became shorter but a few times become longer; and the variance of the times becomes larger.

## 6.2 Definitions of Fractional Derivatives

The various definitions for fractional derivatives exist. A commonly used definition is the Riemann-Liouville fractional derivative [43].

**Definition**: The Riemann-Liouville’s fractional derivative of order *α* is defined as

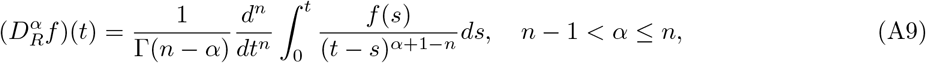

where *α* > 0 is the order of the derivative and *n* ∈ ℕ is the smallest integer greater than *α*. For the Riemann-Liouville’s derivative we have

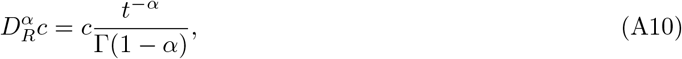

where *c* is a constant. Replacing *c* with *t*^ν^ we get

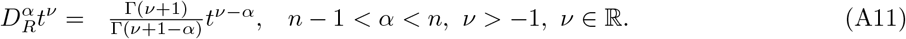

## 6.3 The Mittag-Leffler Functions

The Mittag-Leffler functions 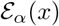 with *α* > 0 have been introduced as [44]

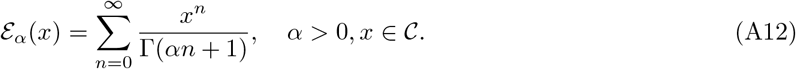

This function provides a simple generalization of the exponential function because of the substitution in the exponential series of *n*! = Γ(*n* + 1) with (*αn*)! = Γ(*αn* + 1). A straightforward generalization of the Mittag-Leffler function is obtained by replacing the additive constant 1 in the argument of the *Gamma* function in Eq. (A12) by an arbitrary complex parameter *β*. For this function we use the following notation

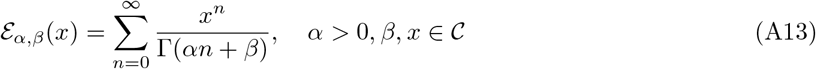

of course 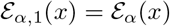. The Mittag-Leffler function can be expressed as a mixture of exponentials [22]

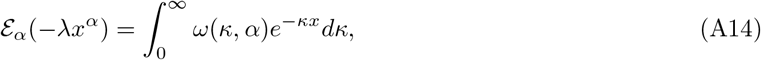

where *ω*(*κ*, *α*) is the probability density

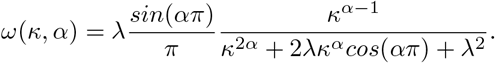

The discrete form of Eq. (A14) is

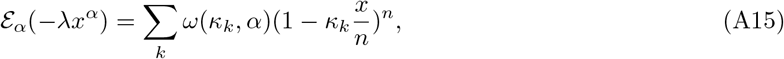

as *k* → ∞. If *n* → ∞ then 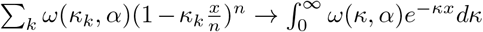. It should be noted since summation in Eq. (A15) is converging as *n* → ∞, we can rewrite Eq. (A15) as

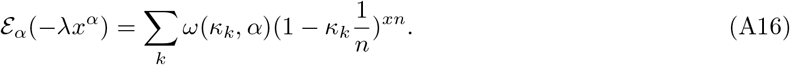

A Mittag-Leffler matrix function [22] can be introduced as continuous form

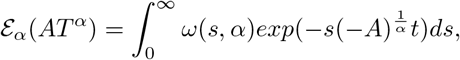

where λ =1 and a discrete form

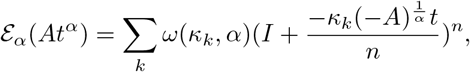

as *k* → ∞ where *A* ∈ *R*^*n* × *n*^ and λ =1.

## 6.4 Proof of theorems related to the TMRCA

In this section, we present the proof of theorems which have been presented on section Time to the most recent common ancestor (TMRCA)

### Theorem 1

**If** 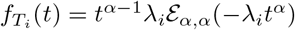 **is the distribution of a waiting time in the** *f*-**coalescent where** *T*_*i*_, *i* = 2, …,*n* **are the coalescent times and** 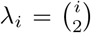 then the distribution of 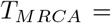 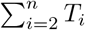 **is as follows**

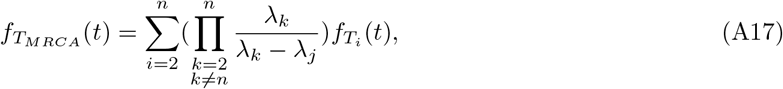

**or, equivalently, this can be presented as**

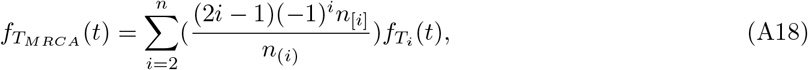

**where**

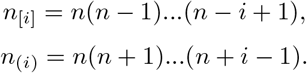

**Proof**: If 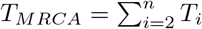 then its distribution is as follows

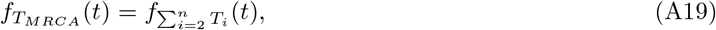

using the Laplace transform we have

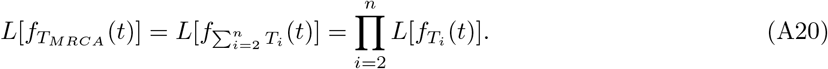

Also using the property of Mittage-Leffler function [45] we know

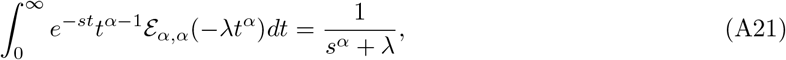

using Eqs. (A20) and (A21) we have

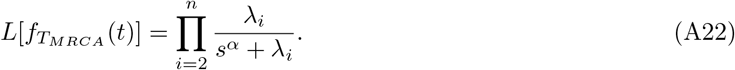

Using Eq. (A22) and partial fraction expansion we have

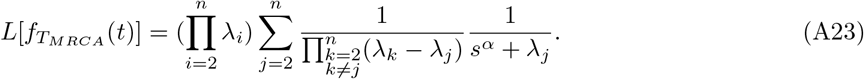

Using Eq. (A23) and Laplace inverse we have

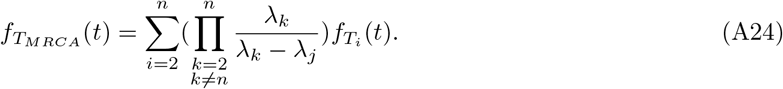

Also we know 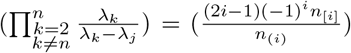 [21] so by using Eqs. (A23) and (A24) we can derive Eq. (A18).

### Theorem 2

**In** *f*-**coalescent, the probability** *P*_*nm*_(*T*) **that** *n* **genes are descendants from** *m* **genes** *T* **units of time ago is**

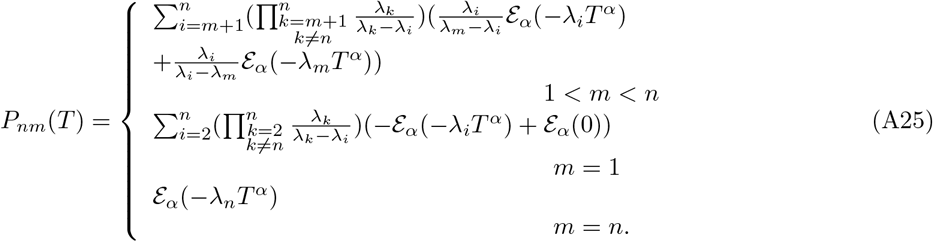

**Proof**: If 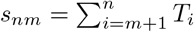 where *T*_*i*_ is the a coalescent time then the probability *P*_*nm*_(*T*) that *n* genes are descended from *m* ancestral genes *T* units of time ago is as follows

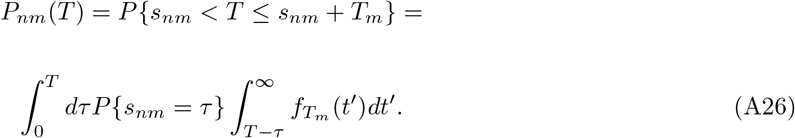

Also we know

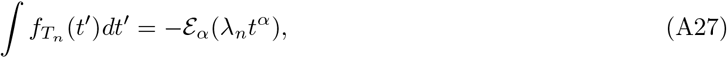

using Eqs. (A17), (A26) and (A27) we have

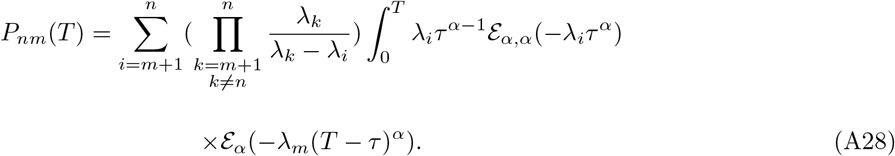

Now we calculate the integral which is in Eq. (A28). If

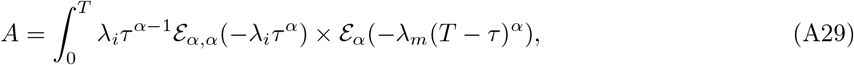

using the Laplace transform and partial fractional expansion we get

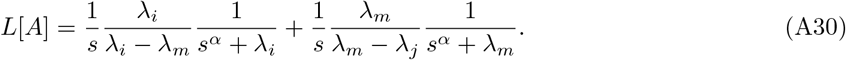

Since 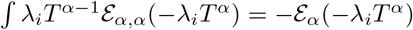, using Eq. (A30) and the Laplace inverse we have

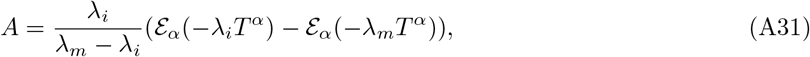

using Eqs. (A28) and (A31) we can derive Eq. (A25) for 1 < *m* < *n*, and we calculate *P*_*n*1_(*T*) and *P*_*nn*_(*T*) as follows

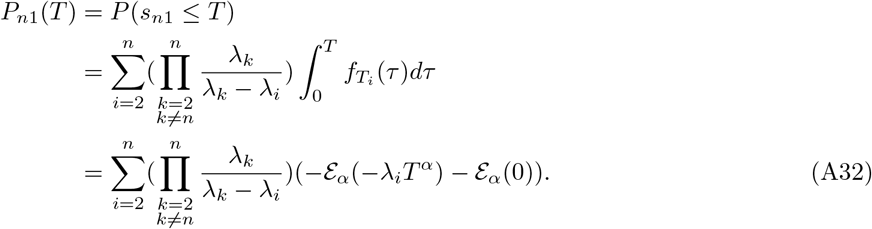

Also we have

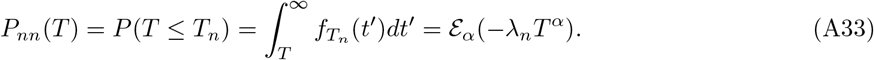

## 7 Supporting Information (SI)

### 7.1 Numerical values of the probability that *n* genes are descended from *m* ancestral

Table 1 gives the numerical values of the probability that *n* genes sampled from a population are descended from *m* ancestral genes *T* units of time ago (*P*_*nm*_ (*t*) in Eq. (24)) for different value of *α*. It is noted that *m* decrease quite rapidly as *T* increase for *n*-coalescent (*α* = 1), but this not the case for *f*-coalescent. The similar results related to *n*-coalescent has been reported in [24]

**Table 1.**
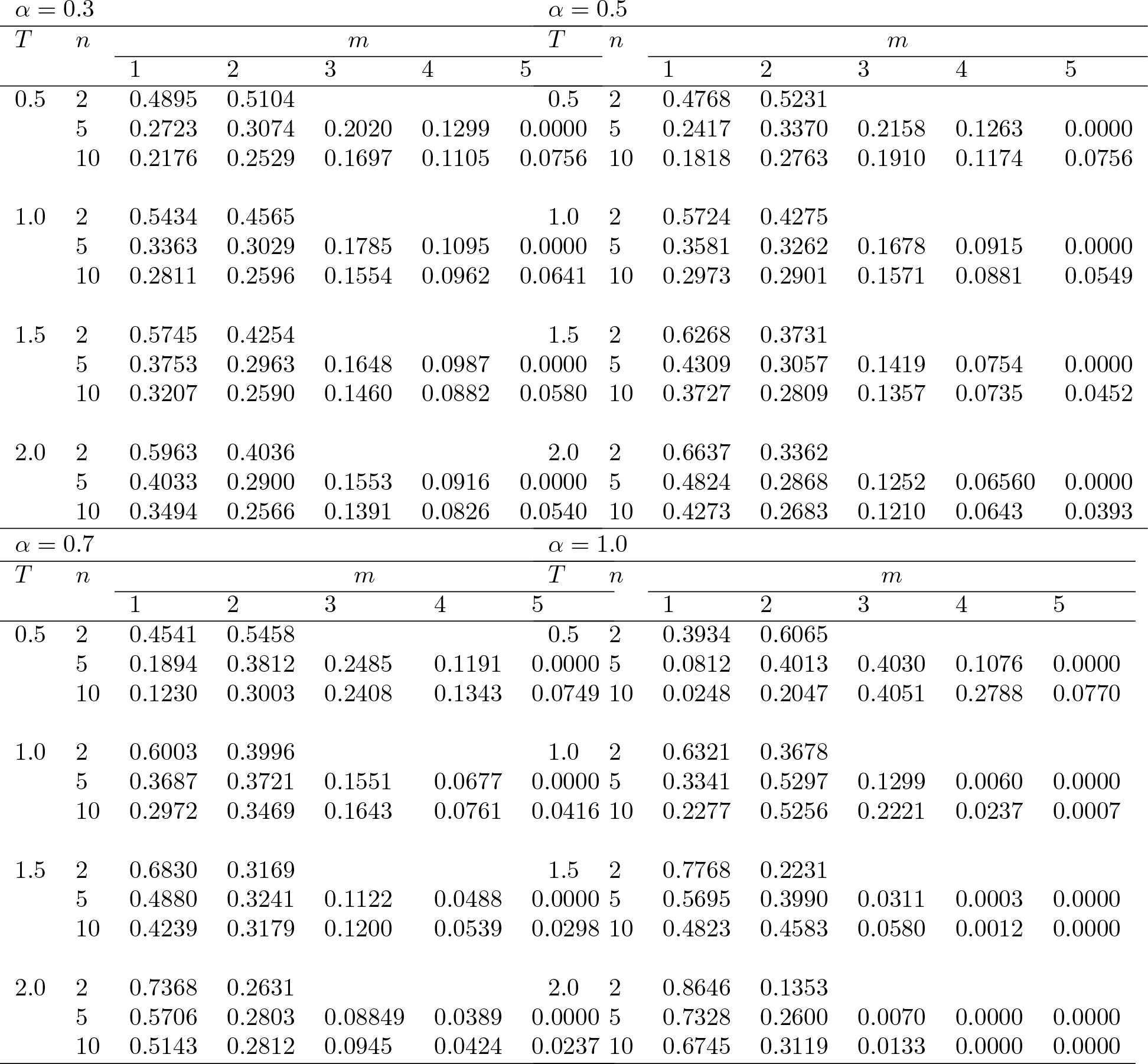
Probability values for *n* genes that are descendants from *m* ancestral genes after time *T* using the *f*-coalescent with different *α* values.

### 7.2 Using the *f*-coalescent for structured coalescent models

The fractional coalescent introduces a parameter that is linked to the variance of waiting times between events. The standard Kingman’s *n*-coalescent has exponentially distributed waiting times that depend on the rate of coalescent and the number of lineages at risk to coalesce. The *f*-coalescent uses the same rate of coalescence and lineages, but allows for further variability; with a low *α* the variability is high and with *α* = 1 the variability is equivalent to the *n*-coalescent. When the population is structured the immigration process can also introduce variability of waiting times between coalescent events. Here we evaluate whether this variability influences *α*, so that *α* could be interpreted as an indicator of missed population structure.

We simulated data from two subpopulations (20 individuals each) for 10 loci (10,000 bp each) using mutation-scaled population size Θ_1_ = Θ_2_ = 0.01 and symmetric immigration rates *M*_1→2_ = *M*_3→1_ = 1, 100, and 10,000. Θ is 4× the effective population size *N*_*e*_ × the mutation rate *μ* per site and generation; *M* is the immigration rate scaled by the mutation rate. The scaled immigration rates are equivalent to the more familiar 4*N_e_m* of 0.01,1.0, and 100. We then ran several models: (1) the correct structured *n*-coalescent model with immigration and two subpopulation sizes; (2) panmictic model using the *n*-coalescent, (3) panmictic model using the *f*-coalescent with *α* < 1. We ran models for *α* = 0.5, 0.06, 0.7, 0.8, and 0.9 (All simulation files are available on https://github.com/pbeerli/fractional_coalescent_data). We compared the different models using Bayesian model comparison [31, 32]. Additionally, we repeated the runs for all models by using only the data of a single population; thus, ignoring the second population. This may lead to intervals between coalescent times that cannot be explained by the *n*-coalescent and would potentially allow a process that can accommodate more variable waiting times such as the *f*-coalescent to better fit the data. The model comparison of the structured datasets rejected any model that uses the *f*-coalescent when using the data from both of the simulated locations. The structured *n*-coalescent was preferred (Table 2) for cases when the data was simulated with low or medium magnitude of gene flow; data simulated with high gene flow is indistinguishable from a panmictic population, thus the single-population model was preferred. When analyzing the data using only a one of the two populations (Table 3), with large gene flow the *n*-coalescent model is preferred; with low immigration the *f*-coalescent model with an *α* = 0.9 is preferred over all other models. Rare immigration will introduce new genetic material into a population which will lead to a waiting time between recorded coalescent events that has very low probability in the *n*-coalescent framework but is normal in the *f*-coalescent framework that allows higher variability of the waiting times.

**Table 2:** Model comparison of datasets simulated under the two-population symmetric structured coalescent with 4 parameters (*θ*1, Θ_2_, *M*_2→1_, and *M*_1→2_) analyzed with different models using the data from both locations.

| (a) |  |  |  |  |  |
| --- | --- | --- | --- | --- | --- |
| Simulated parameter | Model | $\ln mL$ | LBF | Model probability | Rank |
| $\Theta_i = 0.01, M_{j \rightarrow i} = 1$ | structured $n$-coalescent | -315237.51 | 0.00 | 1.00 | 1 |
| | $n$-coalescent | -347952.07 | -32714.56 | 0.00 | 2 |
| | $f$-coalescent $\alpha = 0.9$ | -329491.05 | -14253.54 | 0.00 | 3 |
| | $f$-coalescent $\alpha = 0.8$ | -334463.19 | -19225.68 | 0.00 | 4 |
| | $f$-coalescent $\alpha = 0.7$ | -339960.28 | -24722.77 | 0.00 | 5 |
| | $f$-coalescent $\alpha = 0.6$ | -347409.42 | -32171.91 | 0.00 | 6 |
| | $f$-coalescent $\alpha = 0.5$ | -356376.35 | -41138.84 | 0.00 | 7 |
| $\Theta_i = 0.01, M_{j \rightarrow i} = 100$ | structured $n$-coalescent | -200371.94 | 0.00 | 1.00 | 1 |
| | $n$-coalescent | -200591.59 | -219.65 | 0.00 | 2 |
| | $f$-coalescent $\alpha = 0.9$ | -200891.26 | -519.32 | 0.00 | 3 |
| | $f$-coalescent $\alpha = 0.8$ | -201371.88 | -999.94 | 0.00 | 4 |
| | $f$-coalescent $\alpha = 0.7$ | -202014.66 | -1642.72 | 0.00 | 5 |
| | $f$-coalescent $\alpha = 0.6$ | -202922.34 | -2550.40 | 0.00 | 6 |
| | $f$-coalescent $\alpha = 0.5$ | -204198.38 | -3826.44 | 0.00 | 7 |
| $\Theta_i = 0.01, M_{j \rightarrow i} = 10000$ | structured $n$-coalescent | -194729.51 | -46.41 | 0.00 | 2 |
| | $n$-coalescent | -194683.10 | 0.00 | 1.00 | 1 |
| | $f$-coalescent $\alpha = 0.9$ | -194898.44 | -215.34 | 0.00 | 3 |
| | $f$-coalescent $\alpha = 0.8$ | -195213.06 | -529.96 | 0.00 | 4 |
| | $f$-coalescent $\alpha = 0.7$ | -195735.97 | -1052.87 | 0.00 | 5 |
| | $f$-coalescent $\alpha = 0.6$ | -196419.60 | -1736.50 | 0.00 | 6 |
| | $f$-coalescent $\alpha = 0.5$ | -197584.96 | -2901.86 | 0.00 | 7 |

**Table 3:** Model comparison of datasets simulated under the two-population symmetric structured coalescent with 4 parameters (*θ*1, Θ_2_, *M*_2→1_, and *M*_1→2_) analyzed with different models using the data from the first location.

| Simulated parameter | Model | $\ln mL$ | LBF | Model probability | Rank |
| --- | --- | --- | --- | --- | --- |
| $\Theta_i = 0.01, M_{j \rightarrow i} = 1$ | structured $n$-coalescent | -162693.58 | -236.50 | 0.0000 | 6 |
| | $n$-coalescent | -162493.26 | -36.18 | 0.00 | 3 |
| | $f$-coalescent $\alpha = 0.9$ | -162457.08 | 0.00 | 1.0 | 1 |
| | $f$-coalescent $\alpha = 0.8$ | -162477.44 | -20.36 | 0.00 | 2 |
| | $f$-coalescent $\alpha = 0.7$ | -162551.20 | -94.12 | 0.00 | 4 |
| | $f$-coalescent $\alpha = 0.6$ | -162682.19 | -225.11 | 0.00 | 5 |
| | $f$-coalescent $\alpha = 0.5$ | -162864.37 | -407.29 | 0.00 | 7 |
| $\Theta_i = 0.01, M_{j \rightarrow i} = 100$ | structured $n$-coalescent | -176290.61 | -146.35 | 0.0000 | 3 |
| | $n$-coalescent | -176144.26 | 0.00 | 1.00 | 1 |
| | $f$-coalescent $\alpha = 0.9$ | -176193.29 | -49.03 | 0.00 | 2 |
| | $f$-coalescent $\alpha = 0.8$ | -176294.47 | -150.21 | 0.00 | 4 |
| | $f$-coalescent $\alpha = 0.7$ | -176461.70 | -317.44 | 0.00 | 5 |
| | $f$-coalescent $\alpha = 0.6$ | -176727.94 | -583.68 | 0.00 | 6 |
| | $f$-coalescent $\alpha = 0.5$ | -177088.54 | -944.28 | 0.00 | 7 |
| $\Theta_i = 0.01, M_{j \rightarrow i} = 10000$ | structured $n$-coalescent | -177937.84 | -155.65 | 0.00 | 4 |
| | $n$-coalescent | -177782.19 | 0.00 | 1.00 | 1 |
| | $f$-coalescent $\alpha = 0.9$ | -177814.14 | -31.95 | 0.00 | 2 |
| | $f$-coalescent $\alpha = 0.8$ | -177910.57 | -128.38 | 0.00 | 3 |
| | $f$-coalescent $\alpha = 0.7$ | -178052.81 | -270.62 | 0.00 | 5 |
| | $f$-coalescent $\alpha = 0.6$ | -178291.38 | -509.19 | 0.00 | 6 |
| | $f$-coalescent $\alpha = 0.5$ | -178614.13 | -831.94 | 0.00 | 7 |

We can conclude that the single-population *f*-coalescent is a poor model when the immigration is not very low and when we are aware that there are more populations [cf. 34]. This manuscript discusses the single population *f*-coalescent and we intend to improve on the theory to include multiple populations and include forces such as ongoing immigration and population divergence.

